# SETD2 suppresses tumorigenesis in a KRAS^G12C^-driven lung cancer model and its catalytic activity is regulated by histone acetylation

**DOI:** 10.1101/2025.05.16.654513

**Authors:** Ricardo J. Mack, Natasha M. Flores, Geoffrey C. Fox, Hanyang Dong, Metehan Cebeci, Simone Hausmann, Tourkian Chasan, Jill M. Dowen, Brian D. Strahl, Pawel K. Mazur, Or Gozani

**Author notes:** Equal contribution (Ricardo J. Mack, Natasha M. Flores).

## Abstract

Histone H3 trimethylation at lysine 36 (H3K36me3) is a key chromatin modification that regulates fundamental physiologic and pathologic processes. In humans, SETD2 is the only known enzyme that catalyzes H3K36me3 in somatic cells and is implicated in tumor suppression across multiple cancer types. While there is considerable crosstalk between the SETD2-H3K36me3 axis and other epigenetic modifications, much remains to be understood. Here, we show that SETD2 functions as a potent tumor suppressor in a KRAS^G12C^-driven lung adenocarcinoma (LUAD) mouse model, and that acetylation enhances SETD2 *in vitro* methylation of H3K36 on nucleosome substrates. *In vivo*, *SETD2* ablation accelerates lethality in an autochthonous KRAS^G12C^-driven LUAD mouse tumor model. Biochemical analyses reveal that polyacetylation of histone tails in a nucleosome context promote H3K36 methylation by SETD2. In addition, monoacetylation exerts position-specific effects to stimulate SETD2 methylation activity. In contrast, mono-ubiquitination at various histone sites, including at H2AK119 and H2BK120, does not affect SETD2 methylation of nucleosomes. Together, these findings provide insight into how SETD2 integrates histone modification signals to regulate H3K36 methylation and highlights the potential role of SETD2-associated epigenetic crosstalk in cancer pathogenesis.

## INTRODUCTION

Protein lysine methylation is a common post-translational modification (PTM) that occurs in three distinct states—monomethylation (Kme1), dimethylation (Kme2), and trimethylation (Kme3)—depending on whether one, two or three methyl groups are added to the lysine sidechain (1). Lysine methylation is catalyzed by a class of enzymes named protein <u>lysine m</u>ethyltransferases (KMTs) and removed by protein lysine demethylases (1). Methylation at H3K36 is an evolutionarily conserved histone modification (2). In humans, mutations in the enzymes that determine H3K36 methylation dynamics are linked to a variety of developmental disorders and cancer (2–4). The state and extent of methylation at H3K36 is synthesized by distinct KMTs, with H3K36me2 generated by four related enzymes (NSD1, NSD2, NSD3, and ASH1L), whereas SETD2 is the only human enzyme in somatic cells that has been reproducibly shown to generate H3K36me3 (2, 3, 5). SETD2 and its cognate mark H3K36me3 generally occupy transcriptionally active regions. Functionally, SETD2 influences core molecular processes such as DNA methylation, RNA processing, DNA repair, and genomic integrity (6–11). Notably, while H3K36me2-generating enzymes like NSD2 and NSD3 promote oncogenesis when mutated or overexpressed (e.g. (12–17)), SETD2 is a potent tumor suppressor frequently mutated in clear cell renal cell carcinoma (ccRCC) (9, 18) and several other cancers. For example, the *SETD2* gene (along with other important tumor suppressors) is located within chromosome 3p, a genomic region that is reported to show loss of heterozygosity in ∼90-95% of ccRCC tumors (19). In ccRCC pathogenesis, ∼10-20% of tumors acquire SETD2 mutation on the second intact allele (or more rarely acquire mutations on both alleles) resulting in biallelic loss of *SETD2* and H3K36me3 depletion (9, 18, 19). Deletions and/or loss-of-function mutations in SETD2 are also detected recurrently in different types of leukemia and solid tumors including gastroesophageal cancers and lung adenocarcinoma (LUAD), though at lower frequency than seen in ccRCC (10, 20–23).

SETD2 is a large protein with multiple functional regions and domains beyond its ability to catalyze H3K36me3. Thus, the precise role of H3K36me3 in SETD2-associated functions remain unclear. One established function of SETD2-catalyzed H3K36me3 is to allosterically inhibit the ability of the PRC2 complex to synthesize the repressive H3K27me3 chromatin modification (24–29). Further, H3K36me3, via its recognition by the PWWP domain of DNMT3b, promotes targeted DNA methylation (6). However, beyond H3K27me3 and DNA methylation, potential crosstalk between other epigenetic modifications and SETD2-mediated catalysis of H3K36me3 in the regulation of SETD2 function are relatively unexplored. Here, we test the *in vivo* role of SETD2 in suppressing KRAS^G12C^-driven tumorigenesis in LUAD, as G12C is the most common KRAS mutation in this tumor type, and identify potential regulatory roles for histone acetylation, particularly H3K27 acetylation, in promoting *in vitro* methylation of nucleosomes by SETD2.

## RESULTS

### SETD2 deletion accelerates KRAS^G12C^-driven lung cancer pathogenesis in vivo

Several elegant previous *in vivo* studies utilizing either CRISPR-based or gene knockout approaches demonstrated that loss of SETD2 promotes lung adenocarcinoma (LUAD) tumorigenesis in the canonical KRAS^G12D^-driven LUAD mouse tumor model (22, 30–33). As G12C is the most common KRAS oncogenic variant in LUAD (34, 35), we used a recently developed KRAS^G12C^-driven LUAD mouse model (named *K^c^P*) (36) to test whether the additional loss of SETD2 (named *K^c^P;Setd2*) impacted cancer pathogenesis *in vivo* like it does in a G12D oncogenic mutant background (Figs. 1A-C). In this model, expression of the *Kras^G12C^* mutant allele and homozygous deletions of *p53* and *Setd2* are induced by intratracheal lavage of adenovirus expressing Cre recombinase (Ad-Cre) (36). Following viral infection, *K^c^P* mutant mice develop widespread LUAD with 100% penetrance (see schematic, Fig. 1D; (36)). Consistent with the principal catalytic activity of SETD2 being generation of H3K36me3, *SETD2* deletion resulted in loss of H3K36me3 immuno<u>h</u>isto<u>c</u>hemistry (IHC) signal comparing LUAD tissue sections from *K^c^P* and *K^c^P;Setd2* mice (Fig. 1E). Moreover, *SETD2* depletion accelerated tumorigenesis as measured by a modest increase in tumor nodule numbers and a four-fold increase in tumor burden (Figs. 1F-G). Loss of SETD2 also caused an increase in cellular proliferation in tumors, but did not significantly impact apoptosis levels (Figs. 1E, H-I). Finally, SETD2 depletion resulted in a 40% decline in animal overall median survival time (Fig. 1J). Thus, as previously observed with oncogenic KRAS^G12D^-based models (22, 30–33), SETD2 loss accelerates mutant KRAS^G12C^-driven malignancy *in vivo*.

**Fig 1.**
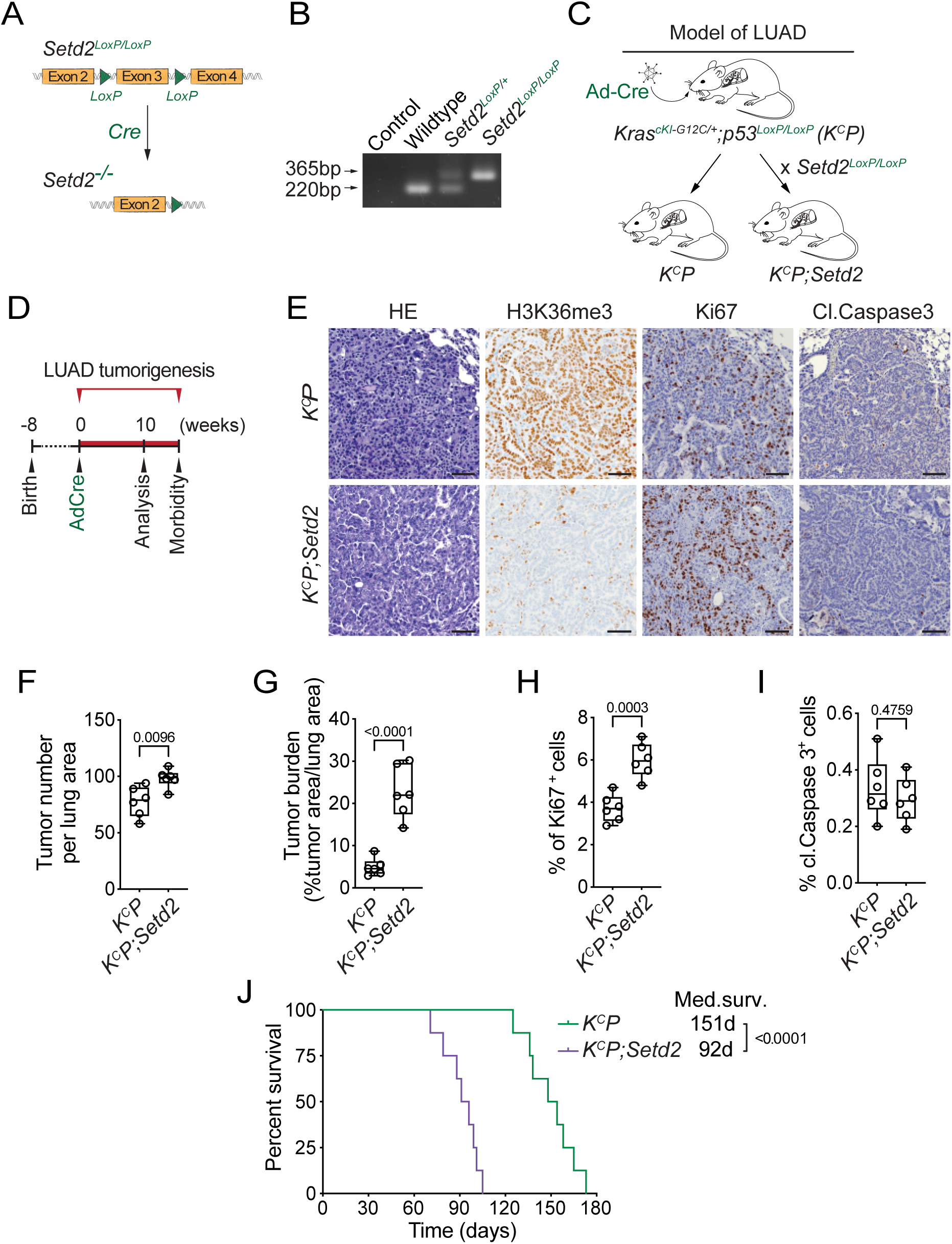
SETD2 ablation promotes KRAS^G12C^-driven lung tumorigenesis in vivo. **A,** Schematic of the *Setd2^LoxP/LoxP^*conditional allele. In the presence of Cre recombinase, exon 3 is deleted to disrupt *Setd2* expression. **B,** Confirmation of *Setd2^LoxP/LoxP^* conditional allele by PCR on DNA isolated from mouse tail biopsies from indicated mouse genotypes, expected product sizes are marked. C, Schematic of generation of LUAD model driven by Cre-recombinase inducible conditional oncogenic *Kras^G12C^* mutation and deletion of p53 (*K^C^P*) and *Setd2* (*K^C^P;Setd2*). **D,** Experimental design to assess effects of SETD2 ablation on LUAD pathogenesis in *K^C^P* model. **E,** Representative HE and IHC staining with indicated antibodies of lung tumors from *K^C^P* and *K^C^P;Setd2* mutant mice at 10 weeks after Ad-Cre induction (n=6/group). H3K36me3 serves as a proxy of SETD2 ablation in tumor cells in *K^C^P;Setd2* mutant mice. P values determined by two-tailed unpaired t-test; boxes: 25th to 75th percentile, whiskers: min. to max., center line: median; scale bars: 100 µm. **F-I,** Quantification of tumor number, tumor burden, proliferation (Ki67+) and cell death (cleaved Caspase3+) in *K^C^P* and *K^C^P;Setd2* samples as in e. **J,** Kaplan-Meier survival curves of *K^C^P* control (n=8, median survival 151 days) and *K^C^P;Setd2* mutant mice (n=8, median survival 92 days) mutant mice. *P* values determined by the log-rank test.

### SETD2 in vitro methylation activity on H3K36 methylated nucleosomes

In humans, SETD2 is the only enzyme in somatic cells that generates the canonical epigenetic modification H3K36me3 (5). In contrast, four KMTs (NSD1, NSD2, NSD3, and ASH1L) generate H3K36me2 (3). Notably, while SETD2 is tumor suppressive (e.g. see Fig. 1), the H3K36me2 KMTs are generally oncogenic (3). While the specific role of SETD2-catalyzed H3K36me3 in cancer pathogenesis remains unclear, it is intriguing that the di-methyl and tri-methyl states at H3K36 may have profoundly different impacts on tumorigenesis. In this regard, it has been postulated that SETD2 requires pre-existing H3K36me2 to generate the tri-methyl state at K36 (2, 3). However, previous work demonstrated that the isolated catalytic SET domain of SETD2 methylates unmodified <u>r</u>ecombinant <u>Nuc</u>leosome<u>s</u> (rNucs) *in vitro* with the same efficiency as methyl-lysine analog (MLA)-based H3K_C_36me2 rNucs (37). While MLA chemistry is generally able to faithfully model native methyl-lysine function (e.g., recognition by reader domains), there are examples where the sulfur moiety compromises functionality (38). Thus, we tested the impact of native methylation installed at H3K36 on the enzymatic activity of the catalytic region of SETD2 that encompasses the SET domain as well as the pre- and post-SET regions (hereafter named SETD2_SET_; Fig. 2A).

**Fig 2.**
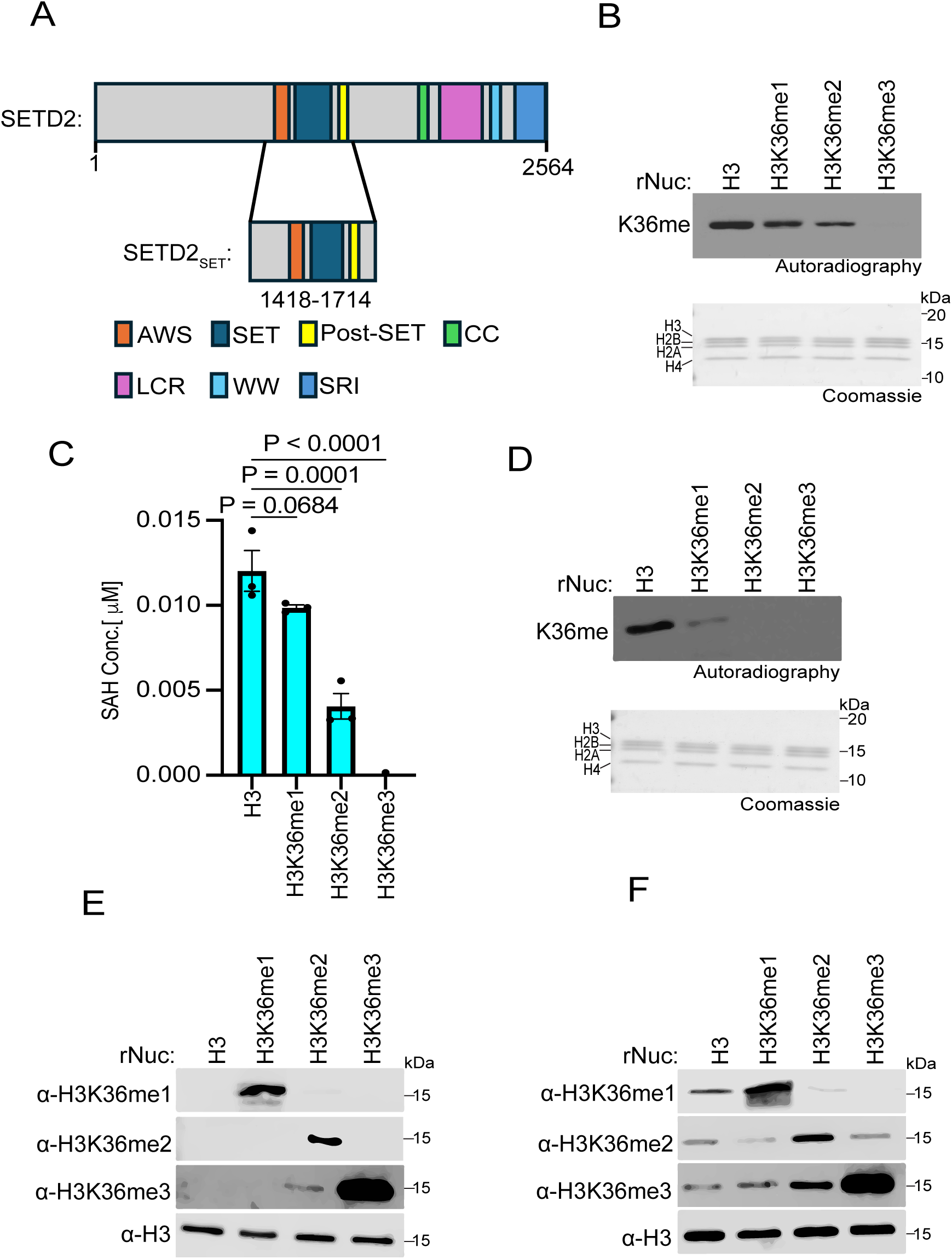
SETD2 methylates unmodified, H3K36me1-, and H3K36me2-modified nucleosomes, but not those bearing H3K36me3. **A,** Schematic of SETD2 domain structure, with the catalytic region used in the biochemical assays (SETD2_SET_) indicated. **B,** SETD2_SET_ *in vitro* methylation reactions with radiolabeled ^3^H-SAM on unmodified (H3), H3K36me1, H3K36me2, or H3K36me3 recombinant nucleosomes (rNuc) substrates as indicated. K36me: methylated H3K36. Top, autoradiography; bottom, Coomassie blue staining. **C,** Methylation (MTase-Glo) assays (see methods) with enzyme and substrates as in (B). SAH concentration serves as a measurement of methylation. Activity is normalized to control conditions. Data are means ± SEM from 3 independent replicates. *P* values determined by one-way ANOVA. **D,** Methylation reactions as in (B) using NSD2_SET_ as the enzyme and the indicated rNucs as substrates. Top, autoradiography; bottom, Coomassie blue staining. **E,** Western blot analysis with the indicated antibodies on the indicated rNucs as in (B). **F,** SETD2_SET_ methylation assays as in (B) using non-radiolabeled SAM and methylation detected by Western analyses using the antibodies characterized in (E). H3 is shown as a loading control.

*In vitro* methylation assays were performed using recombinant SETD2_SET_, radiolabeled *S*- Adenosyl-methionine (SAM) as the methyl donor, and rNucs that were either unmethylated or harboring me1, me2, or me3 at K36 as substrate. The SETD2_SET_ domain methylated all the rNuc substrates besides the tri-methylated sample (Fig. 2B). Specifically, the highest signal was observed using unmodified rNucs, with the intensity of the signal decreasing sequentially on H3K36me1 and H3K36me2 rNuc substrates, likely reflecting the reduced capacity of these H3K36 methylated histones to be further methylated (Fig. 2B). SETD2_SET_ showed no detectable methylation activity on H3K36me3 rNucs, consistent with K36 being 100% saturated with methylation and thus there being no more sites available to be methylated and the specificity of SETD2 for K36 versus other lysine residues on histones (Fig. 2B). Similar results were observed using an independent quantitative method that measures production of *S*-Adenosyl-homocysteine (SAH) – a key byproduct of the methylation reaction (Fig. 2C). Like with SETD2_SET_, the NSD2_SET_ domain showed the strongest methylation activity on unmodified rNucs compared to H3K36me1 rNucs, and NSD2_SET_ has no activity on H3K36me2/3 rNucs, consistent with NSD2 being a selective H3K36me2-KMT (Fig. 2D).

To gain insight into the efficiency of conversion between methyl states at K36 for SETD2, we identified methyl-state specific H3K36 antibodies for the three methyl states (Fig. 2E). We next tested how lysine methyl-state transition on the various H3K36 methylated nucleosome substrates impact SETD2_SET_ activity. To this end, we performed methylation assays with SETD2_SET_ using non-radiolabeled SAM and detected modification of rNucs using the H3K36me state-specific antibodies (Fig. 2E). Under our reaction conditions (see Methods), SETD2_SET_ generated all three methyl states (me1, me2, and me3) when using unmodified rNucs as substrate (Fig. 2F). On H3K36me1-rNucs, SETD2_SET_ generated both higher states of methylation at K36 (H3K36me2 and H3K36me3), whereas H3K36me2-rNucs were naturally only converted to the trimethyl state (Fig. 2F). The conversion to the trimethyl state at K36 was most efficient on H3K36me2-rNucs, which likely reflects the reaction having poor *in vitro* processivity (Fig. 2F). These results indicate that the ability of SETD2_SET_ to generate H3K36me3 *in vitro* does not require pre-existing methylation, consistent with previous studies (37). The data further suggests that, at least *in vitro*, SETD2_SET_ is agnostic about the state of methylation at H3K36 on nucleosome substrates.

### Specific histone acetylation events enhance SETD2 methylation activity

Histone acetylation promotes transcription and other DNA-templated processes through several mechanisms, such as the recognition of acetyl-lysine by reader proteins and the neutralization of DNA-histone interactions (39–41). The combined consequences of such activities decondense chromatin to increase accessibility to the underlying DNA. In addition, neutralizing the positive charge on the lysine side chain facilitates accessibility of KMTs to the unstructured histone tails by disrupting tail-DNA interactions, as was previously shown for the MLL1 KMT complex (40–43). Given the relatively restrictive location of K36 of H3 near the globular domain of the nucleosome, we speculated that histone acetylation might render the residue more accessible to methylation by SETD2. To test this idea, methylation assays with SETD2_SET_ were performed using substrates that were either unmodified rNucs or rNucs with tetra-acetylated lysine residues on the N-terminus tails of H2A, H3, or H4 (see schematic, Fig. 3A). SETD2_SET_ methylation activity as determined by incorporation of radiolabeled SAM or SAH production was enhanced on tetra-acetylated rNucs compared to control rNucs irrespective of the acetylated histone tail (Figs. 3B-C). A similar trend was observed in methylation assays using the NSD2_SET_ domain, though H3 tetra-acetylation seemed to have the biggest impact compared to acetylation of H2A and H4 (Fig. 3D). In contrast, methylation assays using hDOT1L, the KMT responsible for H3K79 methylation – a PTM situated in the nucleosome globular domain – was not enhanced by acetylation of the various histone tails (Fig. 3E). Collectively, these data suggest that a high degree of acetylation may increase the accessibility of H3K36 to the enzymes that methylate this residue.

**Fig 3.**
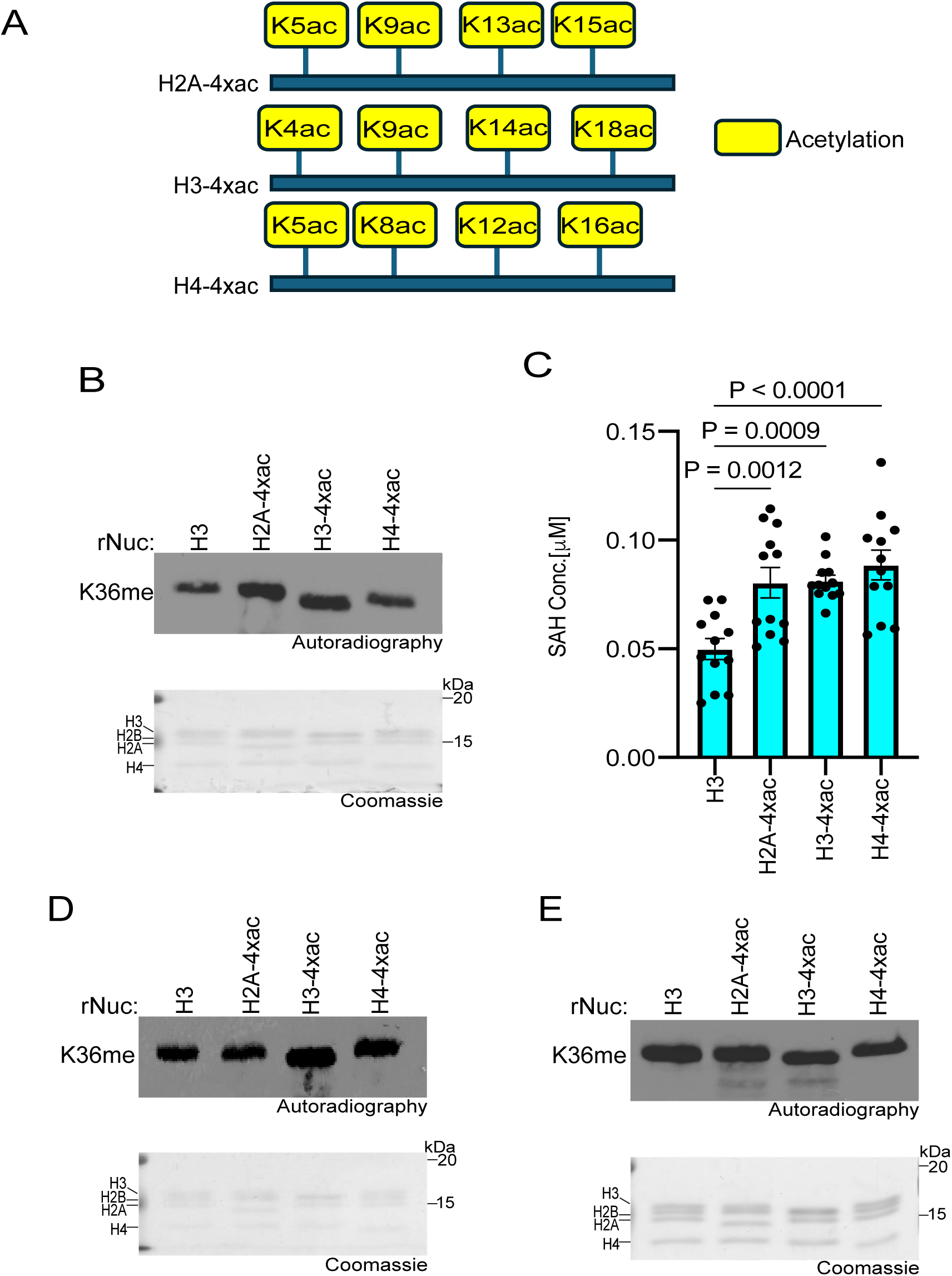
Histone poly-acetylation enhances SETD2_SET_ activity. **A,** Schematic showing the tetra-acetylated rNuc tested in the study. **B,** *In vitro* methylation reactions with SETD2_SET_ as in Fig. 2B using the indicated tetra-acetylated rNucs. Top, autoradiography; bottom, Coomassie blue staining. **C,** MTase-Glo assays as in Fig. 2C using the indicated tetra-acetylated rNucs. Data are means ± SEM from 12 independent replicates. *P* values determined by one-way ANOVA. **D,** Methylation reactions as in (B) using NSD2_SET_ as enzyme. Top, autoradiography; bottom, Coomassie blue staining. **E,** Methylation reactions as in (B) using DOT1L as enzyme. Top, autoradiography; bottom, Coomassie blue staining.

We next explored whether acetylation of individual residues on the H3 tail could facilitate SETD2 activity, and if so, whether such an effect depends on the specific acetylation site. To this end, methylation assays with SETD2_SET_ were performed using substrates that were either unmodified rNucs or rNucs mono-acetylated at K4, K9, K14, K18, K23, or K27 on the N-terminal tail of H3 (see schematic, Fig. 4A). Methylation activity, determined by incorporation of radiolabeled SAM (Fig. 4B) or SAH production (Fig. 4C), indicated that three different single acetylation sites (K14, K23, and K27) enhanced SETD2_SET_ methylation activity on nucleosomes, with the impact likely H3K27ac>H3K14ac>H3K23ac (Figs. 4B-C). While understanding the molecular basis for the acetylation-mediated enhancement in SETD2 activity will likely require structural investigation, we reasoned that given the proximity and relationship between H3K27 and H3K36 (24, 44), that acetylation at K27 might impact the interaction between SETD2_SET_ and nucleosomes. Consistent with this, SETD2_SET_ was subtly more efficiently pulled down when using H3K27ac rNucs as bait compared to unmodified rNucs and the ability of SETD2_SET_ to form a complex with recombinant mono-nucleosomes was subtly enhanced in the presence of K27 acetylation (Figs. 4D-E). These data suggest K27ac may directly or indirectly minorly influence the interaction between SETD2_SET_ and its substrate site in the context of nucleosomes (see Discussion).

**Fig 4.**
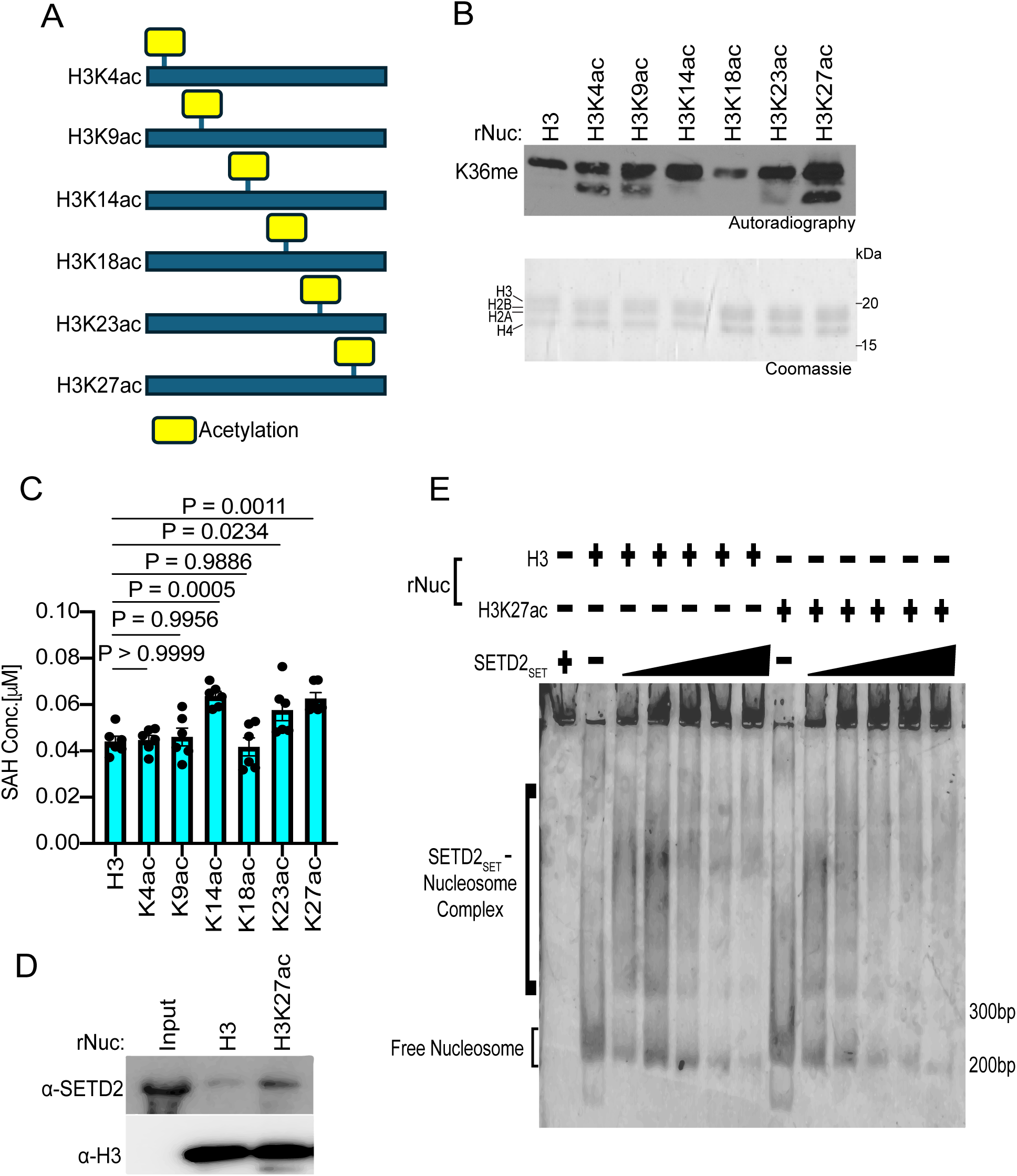
H3K27 acetylation enhances SETD2_SET_ binding and activity. **A,** Schematic showing the different histone H3 acetylated rNuc tested in the study. **B,** *In vitro* methylation reactions with SETD2_SET_ as in Fig. 2B using the indicated acetylated rNucs. Top, autoradiography; bottom, Coomassie blue staining. **C,** MTase-Glo assays as in Fig. 2C using the indicated acetylated rNucs. Data are means ± SEM from 6 independent replicates. *P* values determined by one-way ANOVA. **D,** Nucleosome pulldown assay using biotinylated unmodified or H3K27ac nucleosomes as bait to assess binding to GST-SETD2_SET_. Bound protein was detected by Western blotting with indicated antibodies. **E,** Electrophoretic mobility shift assay (EMSA) with increasing concentrations of SETD2_SET_ (0–5 μM) incubated with the indicated rNuc (250 nM) and SETD2_SET_ binding to rNuc detected by staining DNA with Sybr Gold.

### Histone ubiquitination does not impact SETD2 methylation activity in vitro

We next investigated how ubiquitination at various histone residues influences SETD2 activity. While methylation at H3K36 directly blocks a key allosteric interaction between the PRC2 complex and unmodified H3K36, H3K27me3 indirectly inhibits H3K36 KMTs. Specifically, H3K27me3 recruits the PRC1 complex, which catalyzes the ubiquitination of H2AK119 (H2AK119ub) (45). Nucleosomes harboring H2AK119ub antagonize H3K36 enzymes like NSD2 by structurally preventing their proper association with the nucleosome (44). However, whether H2AK119ub and/or other histone ubiquitination sites impact SETD2 activity is unclear, although studies in yeast have shown the SETD2 homolog, Set2, is influenced by H2BK120ub (46). Thus, we tested SETD2_SET_ activity against a library of ubiquitinated nucleosomes as substrates (see schematic, Fig. 5A). Surprisingly, assaying methylation by three different methods (utilizing radiolabeled SAM, detecting the different methylation states by Western, and measuring SAH generation), histone ubiquitination did not impact SETD2 activity (Fig. 5B-D; note that the H3K36me2 antibody is blocked by H3K14 and H3K18 ubiquitination). In contrast, as expected, H2AK119ub specifically reduced NSD2 activity (Fig. 5E). Collectively, these findings indicate that the isolated catalytic domain of SETD2, unlike for NSD2, is not influenced by histone ubiquitination, suggesting potential functional differences in how H3K36 enzymes interact with the ubiquitinated nucleosome landscape.

**Fig 5.**
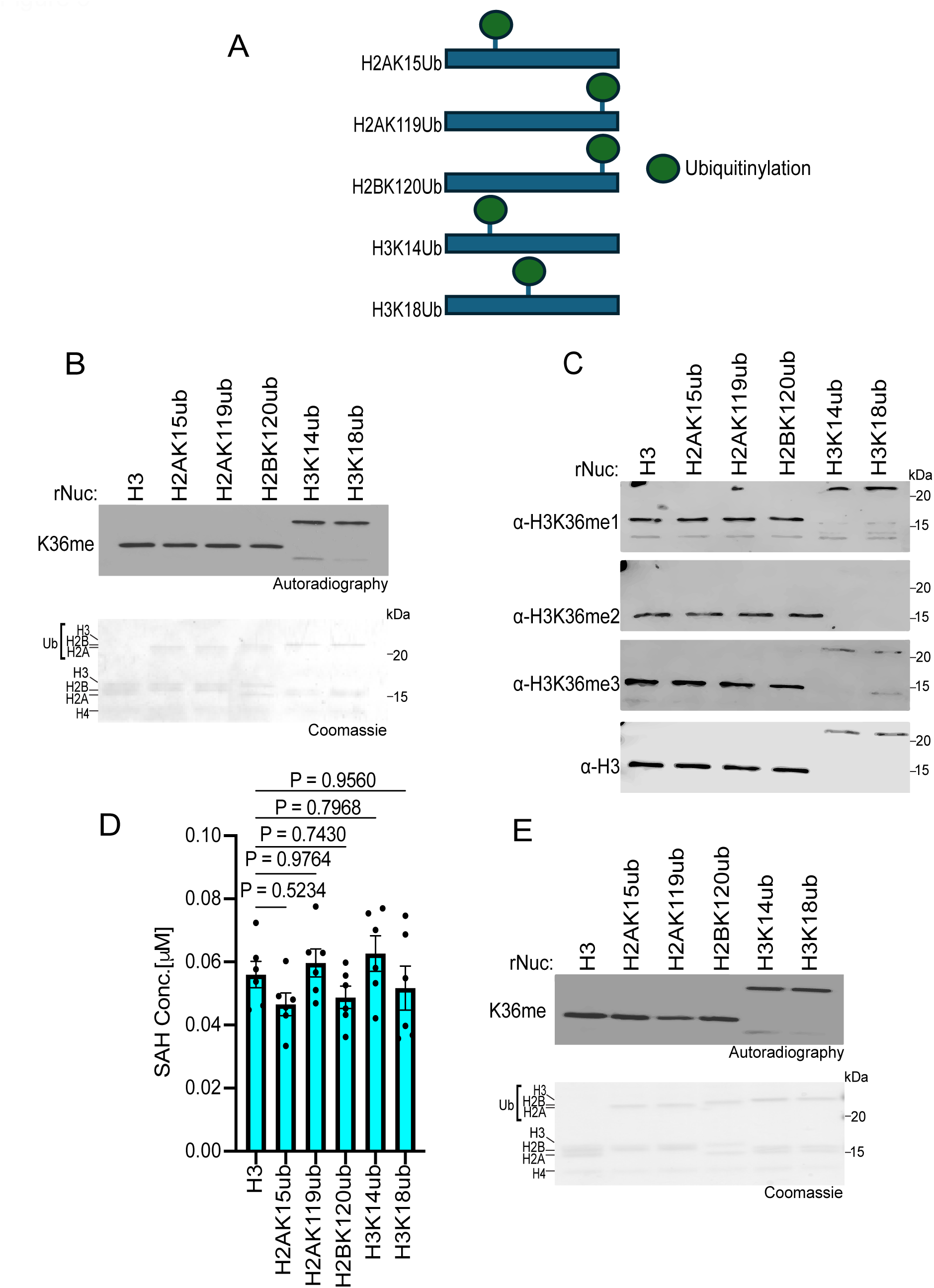
Histone ubiquitination does not affect SETD2_SET_ activity. **A,** Schematic showing the different histone ubiquitinated rNuc tested in the study. **B,** *In vitro* methylation reactions with SETD2_SET_ as in Figure 2B using the indicated ubiquitinated rNucs. Top, autoradiography; bottom, Coomassie blue staining. **C,** SETD2_SET_ methylation assays as in Fig. 2F using non-radiolabeled SAM and methylation detected by Western analysis using the indicated antibodies. H3 is shown as a loading control. Note that H3K14ub and H3K18ub interfere with the αH3K36me2 antibody from recognizing H3K36me2. **D,** MTase-Glo assays as in Fig. 2C using the indicated rNucs. Data are means ± SEM from 6 independent replicates. *P* values determined by one-way ANOVA. **E,** Methylation reactions as in (B) using NSD2_SET_ as enzyme. Top, autoradiography; bottom, Coomassie blue staining.

## DISCUSSION

SETD2, via catalytic and non-catalytic mechanisms, regulates fundamental nuclear processes. Mutations in SETD2 cause Sotos-like syndrome, an overgrowth disorder, and loss of SETD2 commonly occurs and is thought to be tumor suppressive in ccRCC and many other malignancies (3, 47). Indeed, SETD2 was identified as one of the most mutated genes in a saturation analysis of 21 cancer types (48). Consistent with these observations and previous lung cancer mouse modeling studies (22, 32, 33), we find that SETD2 ablation strongly accelerates LUAD malignant progression and lethality in a KRAS^G12C^-driven mouse tumor model (see Fig. 1). While these studies demonstrate that CRISPR-mediated endogenous depletion of SETD2 or homozygous genetic deletion of *SETD2* promote tumorigenesis, future work will help elucidate the specific contributions of H3K36me3 synthesis and other SETD2 functional domains in tumor suppression. In this context, EZM0414, a clinical-grade SETD2 inhibitor, was being evaluated in a Phase I clinical trial as a therapeutic in relapsed or refractory multiple myeloma and diffuse large B cell lymphoma, suggesting that at least in these cancer types, H3K36me3 generation may be oncogenic (NCT05121103; note this trial was terminated). Whether SETD2-mediated catalysis of H3K36me3 is tumor suppressive or oncogenic in a tumor context-dependent manner—and how this activity relates to SETD2’s other functions—are potentially important questions for future investigation.

Consistent with previous work, our biochemical analyses using native designer nucleosomes indicates that—at least in isolation—the catalytic domain of SETD2 does not exhibit a preference for substrates bearing mono- or di-methylation at K36 compared to the unmodified state (see Fig. 2) (37). We speculate that physiologically SETD2 uses all three lower states of methylation at H3K36me (me0-me2) to generate endogenous H3K36me3, with the specific precursor state determined by the underlying biology and genomic context. Our analyses also uncovered an unanticipated regulatory role for histone acetylation in activating SETD2 *in vitro* methylation activity on nucleosomes. Why some acetylation sites increased SETD2 activity whereas others did not is presently unclear. One possibility is that certain acetylation events, for example H3K27ac, may alter histone-DNA interactions in a manner that renders H3K36 more accessible to the catalytic pocket of SETD2. Additionally, our data suggests that SETD2 may interact more strongly with H3K27ac nucleosomes compared to unmodified nucleosomes, which would promote higher levels of methylation (Fig. 4). Finally, we observed a divergence between SETD2 and H3K36me2-KMTs such as NSD2, in that SETD2 is neither inhibited by H2AK119ub nor regulated by H2BK120ub (Fig. 5). These data are consistent with recent cryo-EM-based structural studies, which provide a molecular rationale for why NSD2, but not SETD2, would be impacted by a large modification within the C-terminal region of H2A (44, 49, 50). Future functional and epigenomic studies will be crucial for understanding how epigenetic modifications such as acetylation and ubiquitination regulate SETD2 activity in physiological and pathological settings.

## MATERIALS & METHODS

### Protein expression and purification

The SETD2_SET_ (aa1418-1714, NCBI Sequence: NC_000003.12), NSD2_SET_ (aa959-1365, NCBI Sequence: NC_000004.12), and DOT1L (aa1-416, NCBI Sequence: NC_000019.10) were cloned into pGEX-6P-1 separately. *Escherichia coli* Rossetta cells were transformed with the respective expression vectors and cultured in LB medium (10 g/L tryptone, 5 g/L yeast extract, and 10 g/L NaCl) supplemented with 0.1 mM IPTG (isopropyl 1-thio-ϕ3-D-galactopyranoside, Sigma) at 18 °C for 16–20 h. Cells were lysed using sonicator, lysates were cleared by centrifugation at 12,000 rpm for 20 min and the supernatants were incubated with Glutathione Sepharose (GE Healthcare #17-0756-01); bound proteins were washed and eluted in 10 mM reduced glutathione (Sigma #G4251-25G). Protein concentrations were measured using Pierce Coomassie Plus Assay (Thermo #23236).

For expression and purification of GST-SETD2 (1345–1711) used in Figure 4D, BL21.DE3(pLysS) E. coli transformed with a recombinant expression plasmid encoding GST-tagged human SETD2 catalytic domain (residues 1345-1711) was grown in LB supplemented with 50 µg/mL carbenicillin at 37° C until OD600 reached ∼0.6. Cultures were then transferred to 16° C and induced with 1 mM IPTG (Sigma) overnight and harvested by centrifugation, flash frozen in liquid nitrogen, and stored at −20° C until use. Thawed cell pellets were resuspended in lysis buffer (50 mM HEPES pH 7.5, 150 mM NaCl, 1 mM DTT, 1 mM PMSF) supplemented with 250 U of Pierce Universal Nuclease (ThermoFisher) and 1 mg/mL lysozyme (Sigma) and incubated at 37° C for 10 minutes. Cells were then lysed by sonication (5 x 30 seconds, 40% cycle, 40% power) and lysates clarified by centrifugation before application to glutathione agarose beads (Pierce) pre-equilibrated in wash buffer (50 mM HEPES pH 7.5, 150 mM NaCl, 1 mM DTT, 1 mM PMSF). Following three sequential washes with wash buffer, proteins were eluted in ∼1 mL fractions with wash buffer supplemented with 10 mM glutathione. Fractions containing purified GST-SETD2 were pooled, concentrated by centrifugation filtration (EMD Millipore, MWCO 30 kDa). Glycerol was added to 20% final concentration, aliquoted, and stored at −80°C until use (51).

### In vitro methylation reactions

The methylation reactions on nucleosomes were performed similar as previously described (52). Briefly, 350 nM recombinant enzymes were mixed with 500 nM mononucleosome (EpiCypher, H3-unmodified #16-0006, H3K36me1 #16-0322, H3K36me2 #16-0319, H3K36me3#16-0390, H2A-4xac#16-0376, H3-4xac#16-0336, H4-4xac#16-0313, H3K4ac#16-0342, H3K9ac#16-0314, H3K14ac#16-0343, H3K18ac#16-0372, H3K23ac#16-0364, H3K27ac#16-0365, H2AK15ub#16-0399, H2BK119ub#16-0395, H2AK120ub#16-0396, H3K14ub#16-0398, H3K18ub#16-0401) in reaction buffer (containing 250 mM Tris pH 8.0, 100 mM KCl, 25 mM MgCl_2_ and 50% glycerol), after adding 20 µM SAM (S-adenosylmethionine), the mixture was incubated at 30°C for 3 hours.

### MTase-Glo methyltransferase assay

The MTase-Glo methyltransferase assay kit (Promega #V7602) was used to measure enzymatic activity in the presence of different substrates. Specifically, we established an assay in 10 μL reaction mix containing 35 nM of KMT enzyme, 4 µM SAM, 50 nM mononucleosomes (EpiCypher), and MTase-Glo reagent (1×) in reaction buffer (containing 250 mM Tris pH 8.0, 100 mM KCl, 25 mM MgCl2 and 50% glycerol) arrayed in a white 384-well microplate (Corning #CLS3574). Each independent biochemical reaction was performed in triplicate and incubated for 3 hours at 30°C. Subsequently, 10 μL of MTase-Glo detection solution was added and incubated for 1 hour at room temperature. Reactions were detected by luminescence.

### Western blot analyses

For western blot analysis, protein samples were resolved by SDS–PAGE and transferred to a PVDF membrane. The following antibodies were used (at the indicated dilutions): H3K36me1 (Abclonal #A11141, 1:1000), H3K36me2 (ThermoFisher #701767, 1:1000), H3K36me3 (EpiCypher #13-0058, 1:1,000), H3 (EpiCypher #13-0001, 1:10000).

### Animal models

*Kras^cKI-G12C^* and *Trp53^loxP/loxP^* mutant mice have been described before (36, 53). Conditional *Setd2^loxP/loxP^* mouse strain was obtained from Shanghai Model Organisms Center, Inc (Cat. NO. NM-CKO-190069). Briefly, the Setd2*^loxP/loxP^*targeted knockin sequence includes the Neo-LacZ cassette flanked by Frt sites and exon 3 sequence flanked by LoxP sites. Founder mice were crossed with Rosa26-FlpO deleter strain (54) to generate *Setd2^loxP/loxP^*conditional allele. Confirmation of *Setd2^loxP/loxP^* conditional allele targeting was performed by PCR on DNA isolated from mouse tail biopsies and the following primers: Forward: AGCTGACCTGATTTCTCCTTTAG; Reverse: AACAGCTGAGAGTGACCATGAG. Mice were maintained on a mixed C57BL/6;FVB strain background and we systematically used littermates as controls in all the experiments. Both male and female animals were used in the experiments, and no sex differences were noted. In all experiments, animals were numbered, and experiments were conducted in a blinded fashion. After data collection, genotypes were revealed, and animals were assigned to groups for analysis. None of the mice with the appropriate genotype were excluded from this study or used in any other experiments. All mice were co-housed with littermates (2–5 per cage) in pathogen-free facility with standard controlled temperature of 72 °F, with a humidity of 30–70%, and a light cycle of 12 h on/12 h off set from 7am to 7pm and with unrestricted access to standard food and water under the supervision of veterinarians, in an AALAC-accredited animal facility at the University of Texas M.D. Anderson Cancer Center (MDACC). Mouse handling and care followed the NIH Guide for Care and Use of Laboratory Animals. All animal procedures followed the guidelines of and were approved by the MDACC Institutional Animal Care and Use Committee (IACUC protocol 00001636, PI: Mazur).

To evaluate the effects of SETD2 ablation on the development and progression of lung adenocarcinoma, we utilized *Kras^cKI-G12C/+^*, *Trp53^loxP/loxP^* (*K^C^P*), and *Kras^cKI-G12C/+^*, *Trp53^loxP/loxP^*, *Setd2^loxP/loxP^* (*K^C^P;Setd2*). To generate tumors in the lungs of *K^C^P;Setd2* and control *K^C^P* mutant mice we used replication-deficient adenoviruses expressing Cre-recombinase (Ad-Cre) as previously described (55). Briefly, 8-week-old mice were anesthetized by continuous gaseous infusion of 2% isoflurane for at least 10 min using a veterinary anesthesia system. Ad-Cre was delivered to the lungs by intratracheal lavage. Prior to administration, the virus was precipitated with calcium phosphate to improve the delivery of Cre by increasing the efficiency of viral infection of the lung epithelium. Mice were treated with one dose of 5 × 10^6^ PFU of Ad-Cre. Mice were analyzed for tumor formation and progression at indicated timepoints after viral infection. The survival endpoint was determined by overall health criteria scoring. Mouse health status and weight were checked daily.

### Histology and immunohistochemistry

Tissue specimens were fixed in 4% buffered formalin for 24 hours and stored in 70% ethanol until paraffin embedding. 3-μm sections were stained with hematoxylin and eosin (HE) or used for immunostaining studies. The following antibodies were used (at the indicated dilutions): cleaved Caspase 3 (CST #9664, 1:200), Ki67 (BD Bioscience #550609, 1:1000), H3K36me3 (CST #4909, 1:1000). Immunohistochemistry (IHC) was performed on formalin-fixed, paraffin-embedded tissue (FFPE) sections using a biotin-avidin HRP conjugate method (Vectastain ABC-HRP kit, #PK4000) as described before (56). Sections were developed with DAB and counterstained with hematoxylin. Pictures were taken using a PreciPoint M8 microscope equipped with the PointView software and quantified using Image J software (v1.53k, RRID:SCR_003070) and QuPath (v0.5.1, RRID:SCR_018257).

### Nucleosome pulldown assays

2.5 pmol of GST-SETD2 was added to nucleosome binding buffer (50 mM Tris-Cl pH 7.6, 300 mM NaCl, 0.1% NP-40, 0.5% bovine serum albumin, 10% glycerol) to a final volume of 25 µL. 12.5 pmol of biotinylated nucleosome was added and rotated overnight at 4° C. 1 µL of streptavidin Dynabeads (Fisher) were equilibrated in nucleosome binding buffer and resuspended to a final volume of 7.5 µL. Resuspended beads were then added to the nucleosome-enzyme mixture and rotated at 4° C for 1 hour. Beads were pelleted on a magnet, the supernatant (unbound) fraction was collected, and the beads were washed with 200 µL of nucleosome binding buffer three times. Following the final wash, beads were resuspended in 15 µL of a 1x SDS-PAGE loading dye and stored at 4° C until immunoblotting.

### Electromobility shift assay (EMSA)

Nucleosomes were incubated with recombinant SETD2_SET_ in EMSA buffer (30 mM Tris-HCl pH 7.5, 100 mM KCl, 25 mM MgCl_2_, 6 mM DTT (Dithiothreitol), 0.0075% Tween 20, 12% Glycerol) for 15 min at room temperature and analyzed by native 0.2X TBE-PAGE. Each reaction contained 250 nM of nucleosome with increasing concentrations of domain (0, 1, 2, 3, 4, 5 µM). Gels were stained with Sybr Gold (Thermo #S11494).

### Quantification and statistical analysis

Please refer to the Figure legends or the Experimental Details for description of sample size (n) and statistical details. All values for n are for individual mice or individual samples. Sample sizes were chosen based on previous experience with given experiments. Differences were statistically analyzed by unpaired two-tailed t-test, and one-way ANOVA with Dunnet’s test for multiple comparisons as indicated.

## Acknowledgements

This work was supported in part by grants from the NIH to O.G. (R35 GM139569), O.G. and P.K.M. (R01 CA272844, R01 CA278940), P.K.M. (R01 CA236949, R01 CA266280, R01 CA272843). P.K.M. is also supported by DoD PRCRP Career Development Award (CA181486), CPRIT IIRA (RP220391) and CPRIT Scholar in Cancer Research (RR160078). R.M. is supported by a DARE fellowship. This work was also supported in part by grants from the NIH to B.D.S. (R35 GM126900) and J.M.D. (R35 GM152103). G.C.F. was supported by a predoctoral training grant from the National Institute for General Medical Sciences (T32 GM135128).

## Author Contributions

R.J.M. and N.M.F. contributed equally to this work. They were responsible for experimental design, execution, data analysis, and manuscript preparation for their sections of the work. G.C.F. contributed to analysis of histone acetylation on SETD2 activity and nucleosome binding under the supervision of J.M.D. and B.D.S. H.D. and M.C. helped R.J.M. perform NSD2 assays. S.H. and T.C. contributed to histopathological evaluation and IHC analysis. O.G. and P.K.M. were equally responsible for supervision of research, data interpretation and manuscript preparation.

## Competing Interests

O.G. is a co-scientific founder and shareholders of K36 Therapeutics, Inc. and Alternative Bio, Inc. P.K.M. is a consultant and stockholder of Ikena Oncology, Inc. and Alternative bio, Inc. O.G. and B.D.S. are board members, co-scientific founders, and shareholders of EpiCypher, Inc. The other authors declare no competing interests.

## DATA AVAILABILITY

Plasmids, antibodies, and cell lines generated in this study will be available from the lead contact upon request with a completed material transfer agreement.

## Notes

### Summary of Updates

Revisions added to address comments from Reviewers at eLife.

